# Comprehensive and scalable quantification of splicing differences with MntJULiP

**DOI:** 10.1101/2020.10.26.355941

**Authors:** Guangyu Yang, Sarven Sabunciyan, Liliana Florea

## Abstract

Alternative splicing of mRNA is an essential gene regulatory mechanism with important roles in development and disease. We present MntJULiP, a method for comprehensive and accurate quantification of splicing differences between two or more conditions. MntJULiP implements novel Dirichlet-multinomial and zero-inflated negative binomial models within a Bayesian framework to detect both changes in splicing ratios and in absolute splicing levels of introns with high accuracy, and can find classes of variation overlooked by reference tools. Additionally, a mixture model allows multiple conditions to be compared simultaneously. Highly scalable, it processed hundreds of GTEx samples in <1 hour to reveal splicing constituents of tissue differentiation.

## Introduction

Gene alternative splicing is a fundamental biological process that gives rise to a wide array of protein isoforms with modified properties in plant and animal and plant systems. More than 95% of human genes are alternatively spliced, and high levels were reported in virtually all sequenced eukaryotic species. Most splicing variations are tissue specific, but splicing is also altered by external stimuli^1^ and aberrant splicing has been associated with diseases^2^. Therefore, there is a great need to accurately map and quantify gene splice variants, as well as to identify differences in splicing between conditions.

Current methods aim to detect and quantify alternative splicing from RNA sequencing (RNA-seq) data at the level of transcripts (isoforms), splicing events (exon skipping, mutually exclusive exons, alternative exon ends, intron retention), or primitive features (subexons, introns). Isoform-level quantification methods (Cuffdiff, Cuffdiff2, MISO ^3–5^) require a reference annotation or a reconstructed set of transcripts, and their performance suffers from incompleteness and inaccuracies in the assemblies. Event level methods (DiffSplice, rMATS ^6,7^) are less affected by assembly errors, but represent only a subset of alternative splicing variations. For both of these methods, quantification is further complicated by the ambiguity in assigning reads that map to multiple locations in the genome and multiple transcripts of a gene. In contrast, more recent methods (LeafCutter, MAJIQ, JunctionSeq ^8–10^) target introns, which can be more reliably identified from read alignments, capture a wider variety of splicing variations, and are less ambiguous to quantify, as intron-spanning reads associate with unique gene splice patterns. Methods further differ in how they define splicing differences. Most methods quantify changes in the relative splicing levels of the target feature within a group of mutually exclusive local splicing patterns (LeafCutter, MAJIQ, rMATS, DiffSplice), or alternately identify features with splicing usage inconsistent with the rest of the gene (JunctionSeq, DEXseq^11^). Yet others look for changes in the overall (absolute) abundance levels, as a means to identify changes in isoform regulation leading to functional effects^3,4,12^. Lastly, to increase accuracy, some methods rely on a pre-existing set of gene annotations to identify relevant splicing variations, limiting discovery of novel and potentially condition-specific features. The rich spectrum of methods for alternative splicing quantification and differential analysis offer a multifaceted yet inconsistent view of alternative splicing variation ^13^.

We introduce MntJULiP, a statistical learning method based on a novel mixture Bayesian framework, for detecting differences in splicing between large collections of RNA-seq samples. MntJULiP represents splicing variation at intron level, thus capturing most splicing variations while avoiding the pitfalls of assembly. It infers intron annotations directly from the alignments, making it possible to discover new unannotated candidate markers. MntJULiP detects both differences in intron splicing levels, herein called *differential splicing abundance* (DSA), and differences in intron splicing ratios relative to the local gene output, termed *differential splicing ratio* (DSR). Salient features of MntJULiP include:

i. a novel statistical framework, including a zero-inflated negative binomial mixture model for individual introns, in the DSA model, and a Dirichlet multinomial mixture model for groups of alternatively spliced introns, in the DSR model;
ii. it captures significantly more alternative splicing variation, and more types of variation, than existing tools;
iii. superior performance compared to reference methods, including increased sensitivity in control experiments, and high reproducibility and reduced false positives in comparisons on real data;
iv. a unique mixture model that allows comparison of multiple conditions simultaneously, to aptly capture global variation in complex and time-series experiments; and
v. highly scalable, it could process hundreds of GTEx samples in less than half an hour.

MntJULiP is implemented in Python and is distributed free of charge under a GPL license from https://github.com/splicebox/MntJulip/.

## Results

We assess the performance of all programs on simulated and real RNA-seq data, with varying degrees of splice variation and different dataset sizes. We illustrate MntJULiP’s ability to detect more types of alternative splicing variation in the comparison of hippocampus samples from healthy and epileptic mice. We then demonstrate MntJULiP’s unique capability for simultaneous multi-condition comparisons in a 7-point time series experiment on differentiating mouse taste organoids, and its ability to handle large data sets on a large collection of RNA-seq samples from four human tissues obtained from the GTEx project. We include in the comparisons, as feasible, the state-of-the-art intron-based tools LeafCutter, MAJIQ, JunctionSeq and the event-based rMATS, and Cuffdiff2 as the only tool among them compatible with the DSA test.

### Performance evaluation on simulated data

In a first, controlled experiment we used simulated data, namely 25 control and 25 perturbed samples, to evaluate MntJULiP (DSR), MAJIQ, LeafCutter, and rMATS in detecting differences in splicing ratios, and MntJULiP (DSA) and Cuffdiff2 in detecting differences in splicing abundance (see **Methods** and **Figure 1A**). On the DSR experiment, MntJULiP(DSR) achieved sensitivity 74.5%, which was 8.0-60.0% higher than its competitors, at very high and comparable precision, 97.4%. Notably, Cuffdiff2, which was not designed as a DSR method, had the highest sensitivity at 94.9%, however at a very significant drop in precision, to 46.4%. On the DSA experiment, MntJULiP(DSA) had very high 97.9% sensitivity and 95.3% precision, to Cuffdiff2’s values of 95.9% and 70.3%, respectively. Sensitivity of MntJULiP’s DSA test was also significantly higher than any of the DSR programs’, which ranged between 31.7-50.3%, illustrating the fact that methods developed for DSR detection are in general not suitable to detect changes in splicing abundance. We further examined in more details the programs’ results by gene class. While true positives for all programs were fairly uniformly distributed across the constituent gene categories, false positives for MAJIQ, rMATS and Cuffdiff2 were dominated by genes outside of the simulated gene set, underscoring the difficulty for these programs to effectively distinguish and filter paralogs and other alignment and assembly artifacts (**Supplementary Figure S1** and **Source Data**).

**Figure 1.**
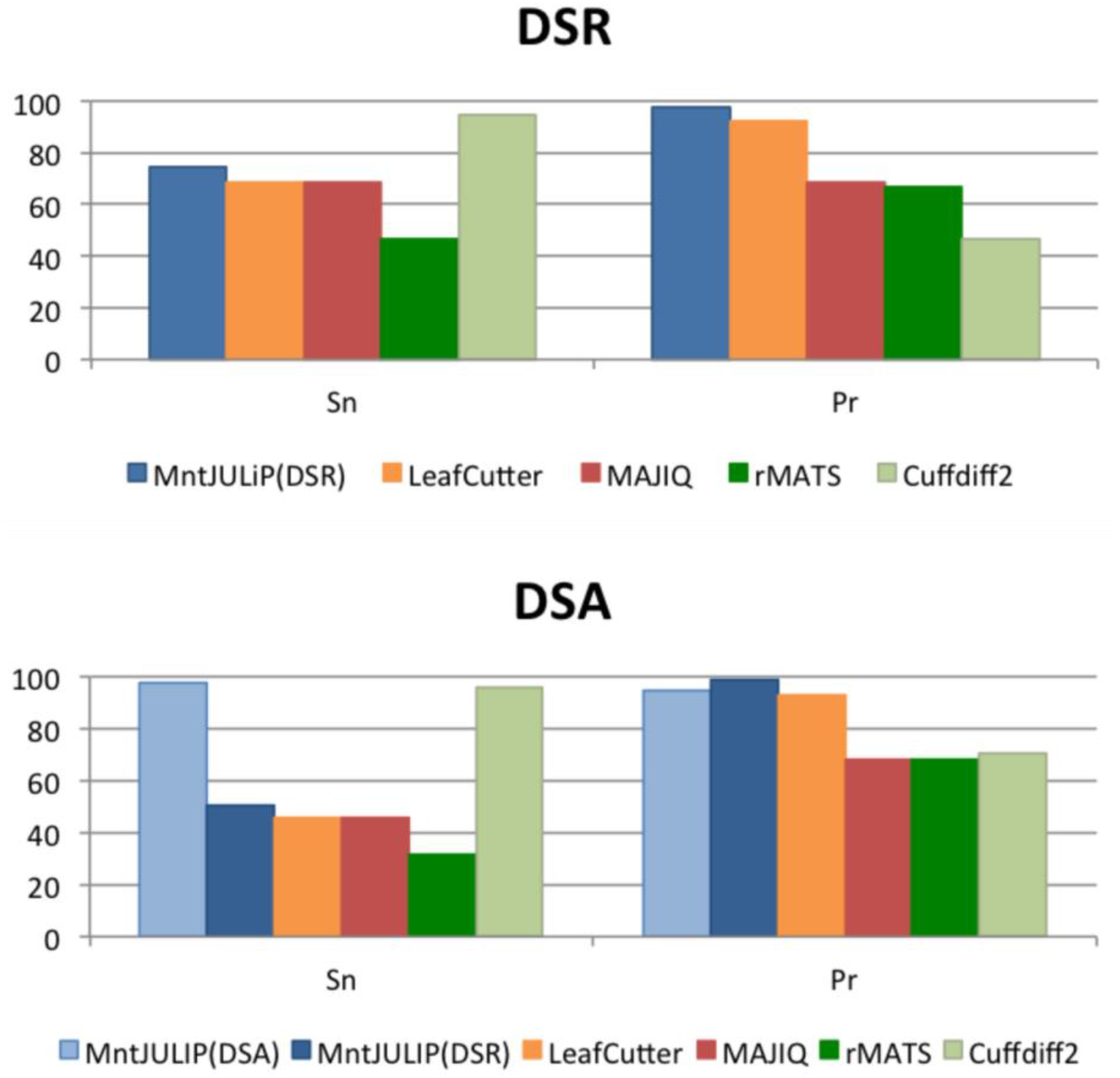
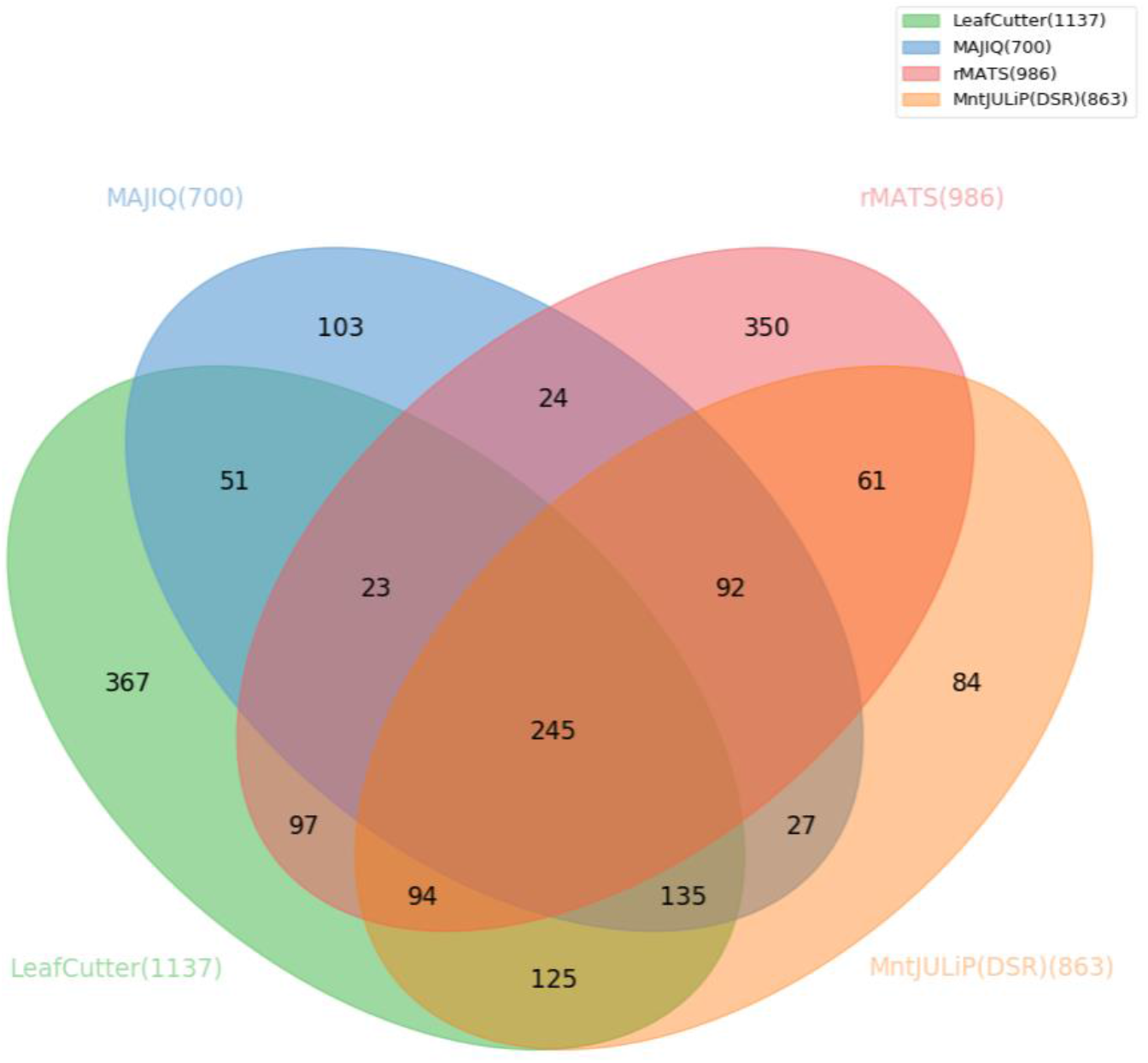
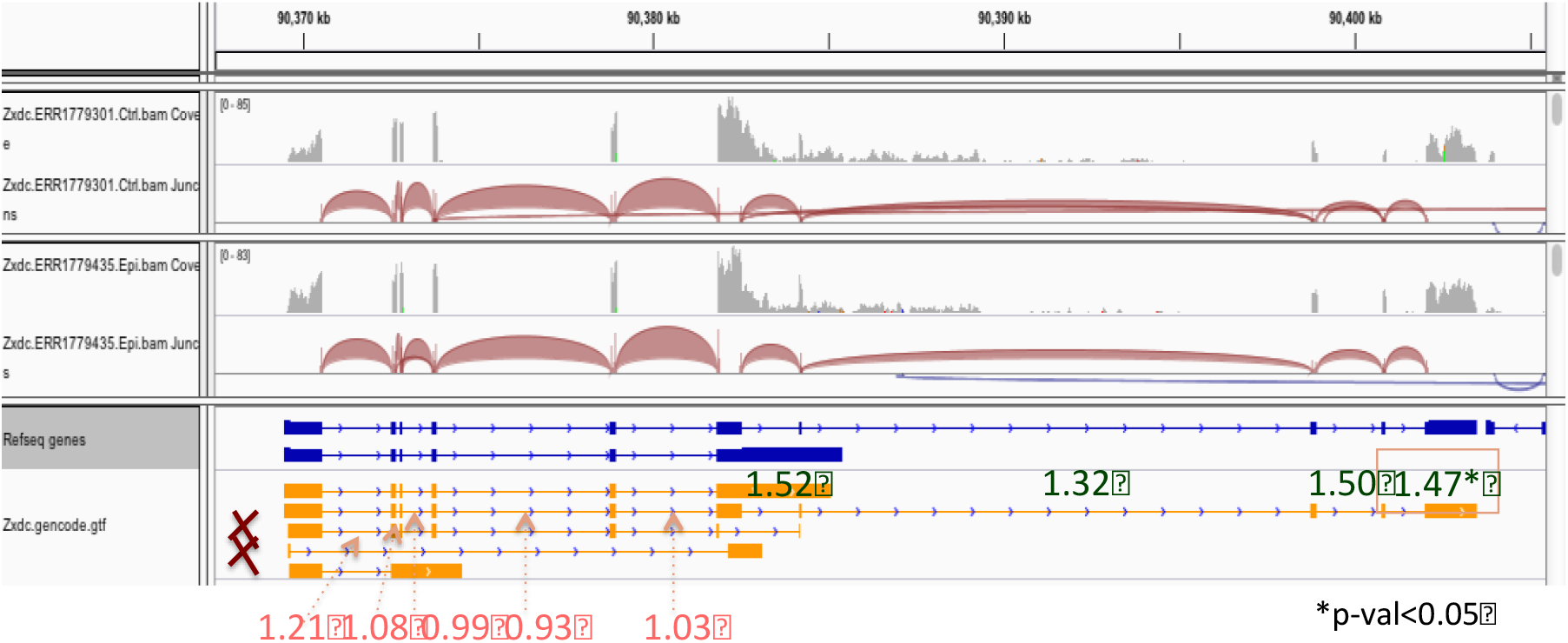
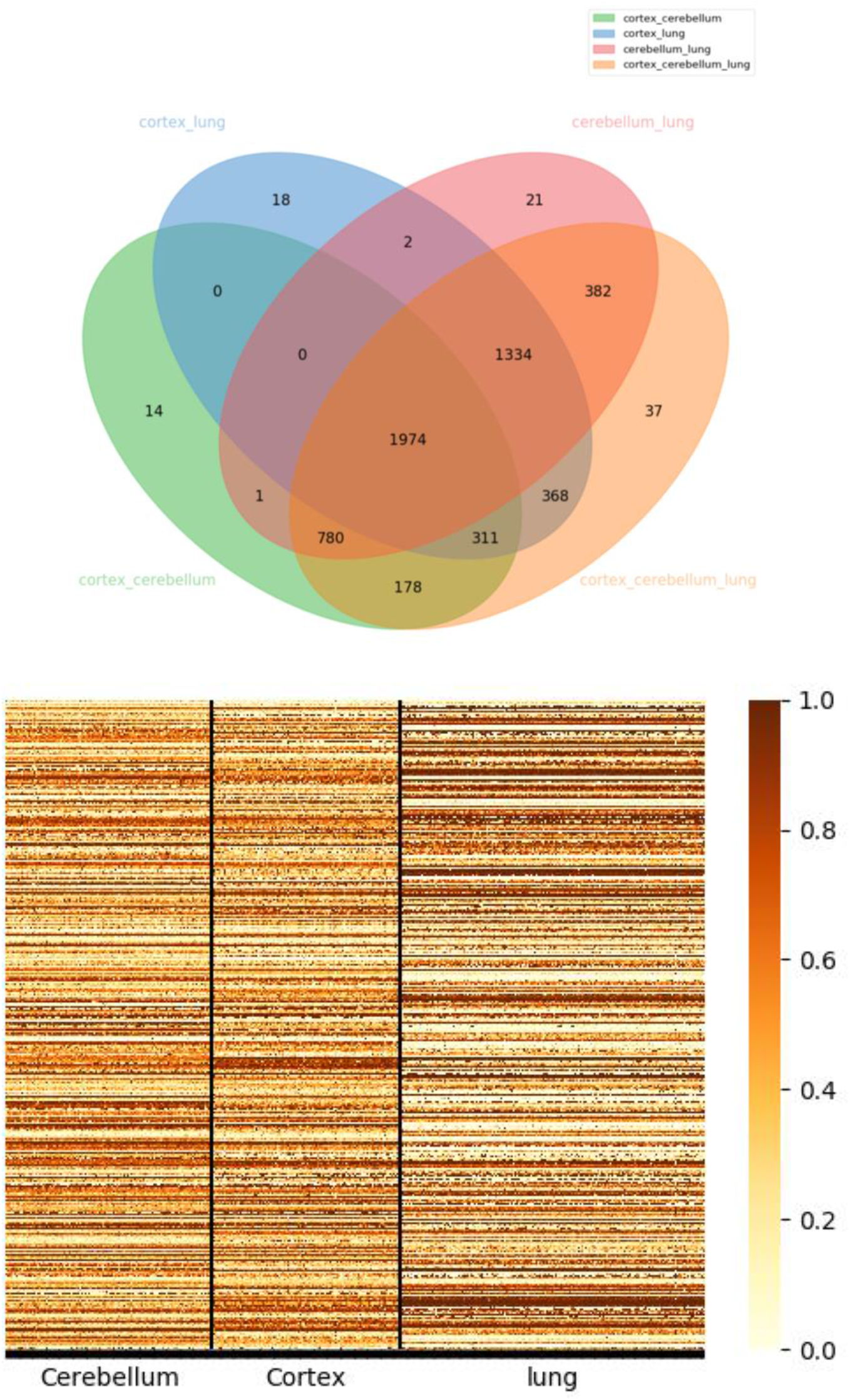
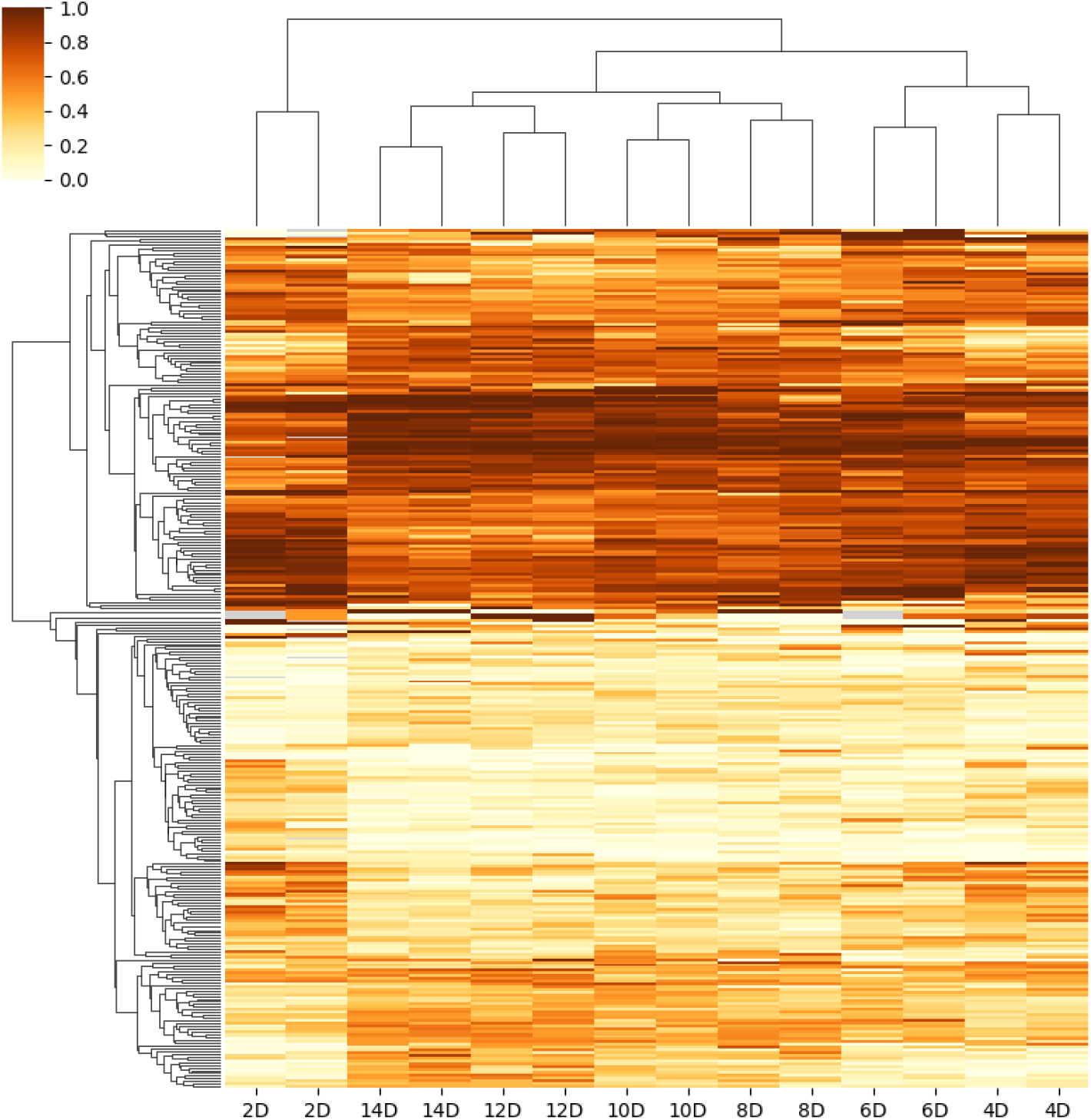
Performance evaluation of MntJULiP on simulated and real data. (A) Comparative evaluation of several methods on 25 control and 25 perturbed simulated RNA-seq data sets. (B) Venn diagram of methods’ gene-level DSR predictions on 24 healthy and 20 epileptic mice. (C) Differential splicing at the *Zxdc* gene locus discovered in the mouse hippocampus data by MntJULiP(DSA); no two introns share an endpoint, therefore the gene could not have been discovered by other tools. Introns are annotated with the fold change values in the comparison of healthy and epileptic mice. (D) Venn diagram of DSR genes, and heatmap of DSR introns discovered with MntJULiP in a multi-way comparison of cerebellum, cortex and lung GTEx RNA-seq samples. (E) Heatmap of DSR introns discovered from the multi-way comparison of 7-stage taste organoid RNA-seq data. Heatmaps show PSI values of differentially spliced introns. Clustering was performed with the Bray-Curtis distance and simple averaging.

We further assessed the methods’ accuracy in quantifying the amount of change in splicing of individual introns (**Supplementary Figure S2**). For the DSR experiment, MntJULiP predictions most closely aligned with the reference annotation (*R*^2^=0.935, Pearson correlation coefficient) between predicted and reference dPSI values, compared to 0.879 for LeafCutter and 0.847 for MAJIQ. For the DSA experiment, MntJULiP had the higher correlation (0.991 versus 0.848) between predicted and reference log fold change values of the two methods. Therefore, MntJULiP predicted values are strongly indicative of the amount of change, and can be used reliably to inform event selection, for instance to select candidate events for experimental validation.

### Performance evaluation on real data

We next applied the methods to RNA-seq samples from hippocampus tissue of 24 healthy mice and 20 mice with pilocarpine induced epilepsy, illustrating a typical RNA-seq experiment. Programs MntJULiP(DSR), LeafCutter, MAJIQ and rMATS predicted between 700 and 1,137 DSR genes (**Figure 1B**). While it is not possible to precisely measure the prediction accuracy in the absence of a ground truth reference, we deem genes predicted by multiple tools as being more reliable. A majority of DSR genes (974 out of 1,878) were predicted by two or more tools. Importantly, MntJULiP had the smallest number and proportion of uniquely predicted genes, 84 (9.7% of its predictions), compared to 350 genes (35.5%) for rMATS, 367 genes (32.3%) for LeafCutter and 103 genes (14.7%) for MAJIQ, and therefore potentially reported the smallest number of putative false positives.

DSR tests capture only a fraction of the alternative splicing variation in an experiment. To showcase the potential of MntJULiP to expand upon the classes of alternative splicing events detected, we assessed the outcomes of MntJULiP’s DSA test. Of the 4,187 genes predicted, 485 were also reported by MntJULiP’s DSR test and an additional 379 by other tools, representing genes with traditional splicing patterns (**Supplementary Figure S3**). An additional 2,510 genes were determined to be differentially expressed by the DESeq2 ^14^ method, a category that is captured by the DSA test. The remaining 813 genes represent a combination of genes with traditional event patterns that could not have been discovered by other tools and putative complex or non-conventional splicing events.

**Figure 1C** and **Supplementary Figure S4** illustrate some of these cases. The Pyruvate Kinase M 1/2 (*Pkm*) gene has two isoforms resulting from the use of mutually exclusive exons **Supplementary Figure S4A**. *Pkm1* is expressed in the adult stage where it promotes oxidative phosphorylation, whereas *Pkm2* is prevalent during embryogenesis and promotes aerobic glycolysis. Splicing dysregulation at this gene has been identified as an oncogenic driver and passenger factor in brain tumors ^15^. While the difference in the isoforms’ splicing ratio is low (0.05) and may have contributed to being missed by other tools, introns flanking both exons yielded positive MntJULiP DSA tests. Most importantly, MntJULiP can detect classes of events that cannot be detected by other methods. In one example at the CWC22 Spliceosome Associated Protein Homolog (*Cwc22*) gene, the two overlapping and mutually exclusive introns at the 3’ end of the gene do not share an endpoint and therefore could not have been interrogated by other methods (**Supplementary Figure S4B**). Similarly, none of the traditional methods can capture variation that results when one isoform’s intron chain is entirely subsumed by another, where the ‘extension’ introns do not share endpoints with others. The ZXD Family Zinc Finger C (*Zxdc*) gene illustrates this example with its 3’ most terminal introns. The GENCODE annotation for this gene lists five isoforms, of which two can be eliminated based on the fact that their unique introns do not appear in any of the 44 samples. Of the remaining isoforms, two have their intron chains entirely subsumed by the longest isoform. In **Figure 1C**, the distribution and average fold change abundance differs significantly between the shared (average 1.03) and isoform specific (average 1.45) intron sets, which can only be explained by a difference in the proportion of splice isoforms in the gene’s output. Lastly, further case analyses revealed other intriguing scenarios, such as at the *Zfp91-Cntf* gene locus (**Supplementary Figure S4C**). The two genes have in common the only intron in the Ciliary Neurotrophic Factor (*Cntf*) gene (chr19:12.764.380-12,765,281), which shows a significant six-fold increase in abundance in the epileptic mice, whereas all other introns for *Zfp91* show a slight decrease within statistical error. While the event can be at first sight attributed to the differential splicing of *Zfp91,* careful observation of the expressed introns reveals that the sole *Zfp91* isoform containing the intron is present at residual levels or not at all in both conditions. Therefore, the increase in abundance appears to be due to the change in the expression of *Cntf,* which owing to the special sharing of gene structure was missed by DESeq2. *Cntf* is a survival factor for multiple neuronal cell types, and an increase in its levels was shown to be involved in attenuating epilepsy-related brain damage ^16,17^.

True accuracy cannot be assessed in analyses on real data. However, to evaluate robustness and reproducibility in the tools’ predictions as an alternative measure of performance ^9^, we divided and analyzed the data into two sets of 10 healthy and 12 epileptic mouse samples. The graphs in **Supplementary Figure S5** show the scatterplots of the estimated difference in percent splicing inclusion (dPSI) between the two replicated experiments. MntJULiP has the highest correlation between the runs (0.579), followed closely by MAJIQ (0.577) and LeafCutter (0.460), and therefore its results are the most robust with the sample set.

### Performance on large data sets

To demonstrate the scalability of MntJULiP and its unique capability to perform simultaneous multi-way comparisons, we applied it to four tissue datasets (frontal cortex, cortex, cerebellum, and lung; 554 samples total) extracted from the GTEx RNA-seq collection. We performed pairwise comparisons as well as three-way comparisons among tissues. In a first experiment comparing the three brain tissues, the multi-way comparison largely recapitulated the individual pairwise comparisons, detecting 99.0% (1,070) of the 1,081 genes and 11 additional genes (**Supplementary Figure S6A**). The test also revealed highly similar splicing profiles between cortex and frontal cortex, with only one gene differentiating the samples. The robustness of the method was confirmed in a second test, comparing the cortex, cerebellum and lung samples (**Figure 1D** and **Supplementary Figure S6C**). All but 14, 18 and 21 of the genes reported from the three pairwise comparisons were selected by the multi-way test, and 37 genes were unique to the three-way comparison, for a 99.3% (5,324 out of 5,364 predicted genes) recovery rate. **Figure 1D** and **Supplementary Figure S7** show the heatmaps of PSI values for each tissue and comparison, reiterating these observations. Similar results can be observed for the DSA test, where the multi-way comparison discovered 97.1% (15,090 out of 15,491) of all genes detected by pairwise comparisons, and only 36 (0.02%) unique genes among the 15,126 predicted (**Supplementary Figure S6D**). Importantly, the comparisons highlighted thousands of differential splicing events that distinguish among the tissues ^18^. Experiments took between 18-44 minutes per comparison on a 24 CPU Intel processor, thereby demonstrating the ability of MntJULiP to handle large-scale applications.

### Application to complex and time-series experiments

All differential splicing methods to date are designed for comparing two conditions, typically ‘cases’ versus ‘controls’. This simple framework is inadequate and impractical for scenarios that involve time-series or complex multi-condition experiments, which seek to determine features that vary across the full range of conditions. As an illustration, we applied both LeafCutter and MntJULiP to RNA sequencing data from mouse taste organoids) at seven growth stages ^19^ (Accession: DRA005238; two samples each at days 2D, 4D, 6D, 8D, 10D, 12D and 14D, for a total of 14 samples). LeafCutter predicted DSR events in 889 genes and MntJULiP in 3,285 genes when combining the results from all-against-all pairwise analyses. By comparison, MntJULiP’s multi-way test predicted 204 differentially spliced genes across all conditions. While true accuracy cannot be measured, we deem features (genes) reported by multiple comparisons to have higher confidence than those predicted in a single comparison, on the basis that features that are differentiated between two stages will likely show variation in other comparisons involving one of the original conditions. As **Supplementary Figure S8** indicates, the distribution of genes according to the number of comparisons in which they are reported is very similar for the LeafCutter and MntJULiP pairwise protocols, with 31-36% of the genes found in only one comparison, pointing to potentially large numbers of false positives. In contrast, the distribution for MntJULiP multi-way predicted genes follows a Bell curve distribution with the mode at 8 comparisons, which provides a more realistic reflection of the experiment. Therefore, the multi-way comparison more accurately identified differences in splicing across the experimental range.

To further examine the landscape of alternative splicing variation during organoid differentiation, we generated heatmaps of the introns discovered with the MntJULiP all-pairwise and the MntJULiP multi-way comparison methods (**Figure 1E** and **Supplementary Figure S9**). Introns’ PSI values show small variation in splicing between consecutive stages, but clear distinguishing characteristics when comparing across all experimental timepoints. In particular, features detected by the multi-way comparison better distinguish between the organoid growth stages, with a significant inflexion point between early (days 2D-6D) and late development and differentiation into taste cells (days 8D-14D), and facilitate more accurate clustering of samples. Interestingly, the visualizations point to distinguishing features separating stage 2D from the other non-differentiated stages, and the separation becomes even more apparent in the DSA visualizations (**Supplementary Figure S9B**). Importantly, these graphical representations highlight the ability of MntJULiP to detect even mild differences between conditions. We also note the ability of MnJULiP to work with very small numbers of samples per condition, as low as two samples per organoid stage.

## Conclusions

We developed MntJULiP, a novel method that detects and quantifies alternative splicing differences at the level of introns, thus avoiding the pitfalls of short read assembly. A variety of methods for differential splicing analysis are currently available, which differ in their selection of target features, objective functions, and technical approaches, leading to poor consistency among the results they produce ^13^. MntJULiP aims to provide a comprehensive view of alternative splicing variation, by representing it at the most granular level (intron) and by implementing two objective functions, aimed at determining differences in the absolute and relative (ratios) intron splicing levels. In comparisons on simulated and real data, we demonstrated that MntJULiP identifies more alternative splicing variation and more classes of variation than other tools, and across a spectrum of experimental conditions, dataset sizes and degrees of variation.

A unique capability of MntJULiP is its ability to perform multi-way comparisons, which is desirable when characterizing complex time series or multi-condition experiments, to identify a global set of features that distinguish among subgroups or stages.

MntJULIp introduces several technical innovations, including its zero-inflated negative binomial and multinomial Dirichlet models to account for low count genes and splice junctions, and the mixture distributions that allow for modeling multiple conditions, thus facilitating multi-way differential analyses.

Lastly, MntJULiP is highly efficient and scalable, providing an effective platform for comprehensive differential splicing analyses of RNA sequencing data from a wide range of experiments and data collections.

## Methods

### Algorithm overview

MntJULiP consists of two components, a ‘builder’ and a ‘quantifier’. The *builder* extracts the splice junctions (introns) and calculates their supporting read counts from the RNA-seq read alignments, filtering introns with fewer than 3 reads in each sample, as potential sequencing and mapping artifacts. (A second filter that removes introns with weak support within the gene’s context is embedded in the statistical model below.) Individual introns are the input to the DSA analysis. For the DSR analysis, introns that share an endpoint are grouped into ‘bunches’. If a reference gene annotation is provided, both individual introns and bunches are associated with an annotated gene if they share at least one intron coordinate. The *quantifier* subsequently evaluates candidate introns, building a learning model for each intron and bunch and performing two statistical tests: i) a test for change in intron abundance (*DSA*), and ii) a test for change in the splicing level of the intron relative to its ‘bunch’ (*DSR*). For the DSA analysis, MntJULiP uses a mixture zero-inflated negative binomial model to estimate individual introns’ abundance levels from the raw read counts. For DSR, it estimates the relative splicing ratios with a mixture Dirichlet-multinomial distribution. For both models, likelihood ratio tests are used to determine the differential splicing events and generate *p*-values, which are then adjusted for multiple testing using the Benjamini-Hochberg correction. The framework is described in detail below.

### A Bayesian read count model

We use a Bayesian statistical framework to estimate the splicing levels of introns for differential analyses. The framework also provides a second filter for weakly supported introns within the context of the gene, by setting a cutoff value for the estimated read count mean. To start, we assume that the read count *y* of intron *v* in a given sample follows a negative binomial distribution *NB*(*μ, θ*). We consider a loose prior with an empirical 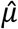 (the sample mean) modeled by a normal distribution: 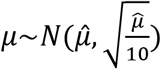 to model the variability between conditions and among the individual samples. Additionally, we apply a restriction on the dispersion parameter with an inverse Half-Cauchy distribution: *φ*^-1^~*HC*(0,5). Lastly, to model low expression introns (0 reads in most samples), we use a zero inflated enhanced negative binomial Bayesian model ^20^:

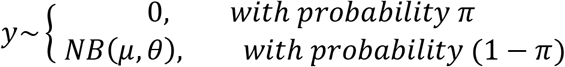

Let *p*(*y*) denote the probability density function for this model. For *n* samples and intron read count *y_j_* in sample *j*, we define the log likelihood:

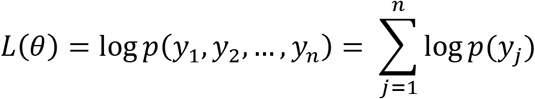

We maximize the log likelihood function using the Limited-memory Broyden–Fletcher–Goldfarb–Shanno (LM-BFGS) optimization method and obtain point estimates for parameters *μ, θ* over the samples.

### The differential splicing abundance (DSA) model

The previous section established the general Bayesian model to estimate intron abundance. Next we describe the framework for modeling individual intron abundance and for DSA testing in a *multi-condition* experiment. Assume that samples are drawn from *m* (typically 2) conditions. Given an intron *ν* and a sample generated from condition *i*, the intron’s read count *y* follows a zero-inflated negative binomial distribution with the condition specific parameters *μ_i_, θ_i_, φ_i_* and *π_i_*, as defined earlier.

Let *p_i_*(*y*) be the probability density function for the complete model for condition *i* = 1… *m*. We define a mixture probability model for *y*.

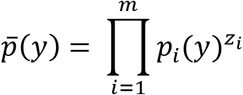

where *z_i_* is the indicator variable for that sample, equal to 1 iff the sample belongs to condition *i* and 0 otherwise.

To formulate the problem, given *n* samples, *m* conditions and *y_j_* the intron read count in sample *j* = 1… *n*, we define the log likelihood:

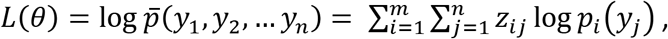

with *Z_ij_* ∈ {0,1} the indicator variable for sample *j* and condition *i*.

Having these two Bayesian models, we establish a hypothesis test for differential intron abundance given the data: the null hypothesis is that samples are generated from the same condition, and the alternative hypothesis is that the samples belong to different conditions, and apply a likelihood-ratio test:

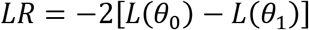

where *L*(*θ_o_*), *L*(*θ*_1_) are the log likelihoods of the null and alternative hypothesis models, respectively, with parameters *θ*_0_ and *θ*_1_.

Lastly, since the parameter *μ_j_* of the alternative hypothesis model is the expected read count (mean) of the intron in condition *j*, we can establish an additional intron filter by setting a threshold for *μ_j_* (e.g., *μ_j_* ≥ 1), to separate a ‘true’ intron from ‘noise’.

### The differential splicing ratio (DSR) model

We next formulate the framework to test for differences in splicing ratios of introns within a ‘bunch’, or group of introns sharing an endpoint. For simplicity, we start by assuming that all samples belong to the same condition and the read counts *y*_1_, *y*_2_,…, *y_k_* in a bundle with *k* introns follow a Dirichlet-multinomial distribution with priors *α*_1_, *α*_2_,…, *α_k_*: *y*_1_, *y*_2_,… *y_k_~DM*(*α*_1_, *α*_2_,…, *α_k_*).

Let *p*(*y*_1_, *y*_2_,…, *y_k_*) be the probability density function of the Dirchlet-mutinomial distribution. For intron read counts *y_j_* = (*y*_1*j*_, *y*_2*j*_,…, *y_kj_*) in sample *j* = 1… *n*, we define the log likelihood function:

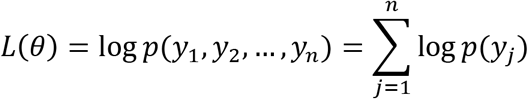

Similar to the discussion in the previous subsection, to extend to the case where samples belong to multiple conditions, we define a Dirichlet-multinomial distribution with prior *α*_*i*1_, *α*_*i*2_,…, *α_ik_* for each condition *i* = 1… *m*:

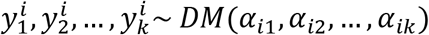

Let *p_i_* = (*y*_1_, *y*_2_,…, *y_k_*) be the probability density function for condition *i*. We define the log likelihood function:

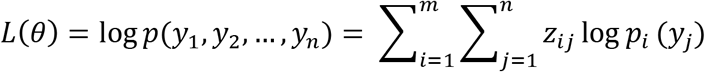

where *y_j_* = (*y*_1*j*_, *y*_2*j*_,…, *y_kj_*) are the read counts of introns in this bunch in sample *j*, *Z_ij_* ∈ {0,1} indicates whether sample *j* belongs to condition *i* or not, and *θ* represents the parameter set of the model.

With the two Bayesian models above, we formulate a log-likelihood ratio test as before: the null hypothesis assumes all samples belong to the same condition, and the alternative hypothesis assumes multiple conditions. Under the alternative hypothesis, the parameters *α*_*i*1_, *α*_*i*2_,…, *α_ik_* for condition *i* can be used to define the splicing ratio, similar to Percent Splicing Inclusion (PSI) ^8,21^, Ψ_*il*_ for intron *I* = 1… *k* under condition *i*, as:

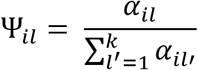

### Sequences and Materials

#### Simulated data

We generated 25 control and 25 perturbed RNA-seq samples with ~86 million 101 bp paired-end reads each, using the software Polyester with human GENCODE v.22 as reference annotation. For the control samples, we used a model of gene and transcript abundance inferred from lung fibroblasts (GenBank Accession: SRR493366). To simulate the perturbed condition, we randomly selected 2,000 annotated protein coding genes with two or more expressed isoforms and assigned them to four groups as follows ^10,22^: i) 500 genes were left unperturbed (NONE); ii) 500 genes had only expression changes (DE), where genes were randomly assigned one half or double the original FPKM value; ii) 500 genes had only splicing differences (DS), obtained by swapping the expression values of the top two isoforms; and iv) 500 genes had both expression and splicing changes (DE-DS). Thus, 1,500 genes underwent changes in splicing abundance, and 1,000 had differences in splicing, and were used as the gold reference for evaluating the tools under the DSA and DSR models, respectively.

#### Real data

Reads for 44 mouse hippocampus samples (24 cases and 20 controls) were obtained from GenBank (Project ID: PRJEB18790). Tissue RNA-seq samples for comparative analyses (121 cortex, 105 frontal cortex, 132 cerebellum, and 196 lung samples) were obtained from the GTEx collection ^23^. Lastly, RNA-seq data from differentiating mouse taste organoids ^19^ (14 samples, 7 stages) were obtained from the Sequence Read Archive (Accession: DRA005238). Listings of the sample IDs are provided in the **Supplementary Source Data**.

### Performance evaluation

Reads were mapped with the program STAR v.2.4.2a ^24^ to the human genome GRCh38 or mouse genome GRCm38 (mm10), as applicable. Alignments were analyzed with the programs MntJULiP v1.0, LeafCutter v0.2.8, MAJIQ v1.1.7a, rMATS v3.2.5 and Cuffdiff2 v2.2.1 to determine changes in alternative splicing profiles. For the simulated tests, transcripts were reconstructed across each sample with StringTie v2.1.4 then merged across samples with StringTie(ST)-merge and the GENCODE transcripts as reference, to create a set of gene annotations to be used with all programs. To evaluate the programs’ accuracy in predicting differentially spliced genes from the simulated data, the 1000 (DS, DE-DS) gene set and the 1,500 (DS, DE, DE-DS) gene set were used as the gold standard for DSR and DSA prediction, respectively. Any other program predictions were deemed false positives. Standard sensitivity (Sn = TP/(TP+FN)), precision (Pr = TP/(TP+SP)), and the F1 = Sn*Pr/(Sn+Pr) value were used to measure accuracy. To assess the programs’ fidelity in quantifying alternative splicing for the DSR test, reference Percent Splice Inclusion (PSI) values for all reference introns were calculated from the simulated data, as the ratio between the intron abundance and that of its bunch. Similarly, for the DSA test, reference log fold change values were calculated for each intron as the log fold change of the cumulative expression levels of all splice isoforms containing that intron.

## Acknowledgements

Development and evaluations were performed on the Maryland Advanced Research Computing Center (MARCC). Work was supported in part by NIH grant R01GM129085 to L.F. and NIH grant R01GM124531 to L.F. and Kathleen Burns. Sarven Sabunciyan was supported by a grant from the Stanley Medical Research Institute.

## Author contributions

L.F. and G.Y. conceived the project, G.Y. developed the method, S.S. provided critical discussions, and G.Y., L.F. and S.S. contributed to the evaluation. All authors wrote and approved the manuscript.

