## Supplemental Figures for "Comprehensive and scalable quantification of splicing differences with MntJULiP"

Table of Contents:

**Supplementary Figures and Tables**

**Figure S1.** Program predictions by gene category in the simulation experiment.
**Figure S2.** Quantification accuracy of programs at intron level in the simulation experiment.
**Figure S3.** Venn diagram of programs' predictions on the mouse hippocampus data set.
**Figure S4.** Examples of MntJULiP predictions not identified with other tools (mouse hippocampus data set).
**Figure S5.** Reproducibility plots for MntJULiP, LeafCutter and MAJIQ (DSR test) and MntJULiP (DSA test) on the mouse hippocampus data.
**Figure S6.** Multi-way versus all-against-all pairwise comparisons on GTEx tissue samples - gene sets.
**Figure S7.** Multi-way versus all-against-all pairwise comparisons on GTEx tissue samples - heatmaps.
**Figure S8.** Continuity-based assessment of program predicted features for the time-series taste organoid data set.
**Figure S9.** Heatmaps of differentially spliced features in the taste organoid data set.

**Figure S1. Program predictions by gene category in the simulation experiment.** Breakdown of programs' predictions by the four gene categories (DS, DE, DE-DS and NONE), and novel (NA), i.e. not in the simulation set.

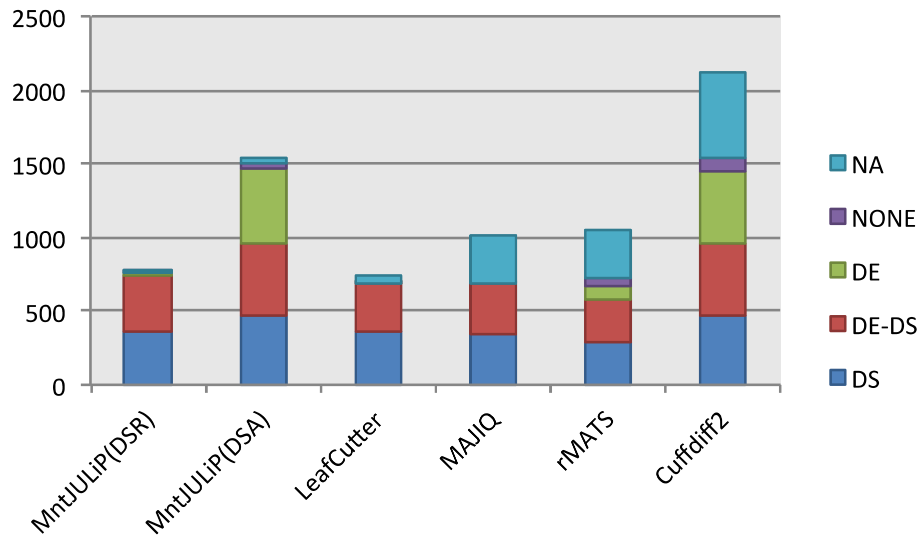

**Figure S2. Quantification accuracy of programs at intron level in the simulation experiment**. (A) _DSR test:_ Scatterplots of reference and predicted dPSI = $PSI_{test}$-$PSI_{ctrl}$ values for 11,282 (MntJULiP), 12,023 (LeafCutter) and 23,406 (MAJIQ) reported introns. Red points mark predictions with p-val<=0.1 and |dPSI|>=0.05. (B) _DSA test:_ Scatterplots of reference and predicted log2fc = $\log_2 (N_{test}/N_{ctrl})$ values for 31,868 reported introns (MntJULiP) and 10,305 (Cuffdiff2) isoforms. The p-val=0.1 line corresponding to the programs' prediction cutoff is marked with a dotted line. (C) Table of Pearson correlation coefficients for the DSR and DSA scatterplots above, considering all introns (isoforms, for Cuffdiff2) evaluated by a program and introns (isoforms) correctly predicted by the program to be differentially spliced (TPs), respectively. MntJULiP shows the highest correlation in each category.

(A)

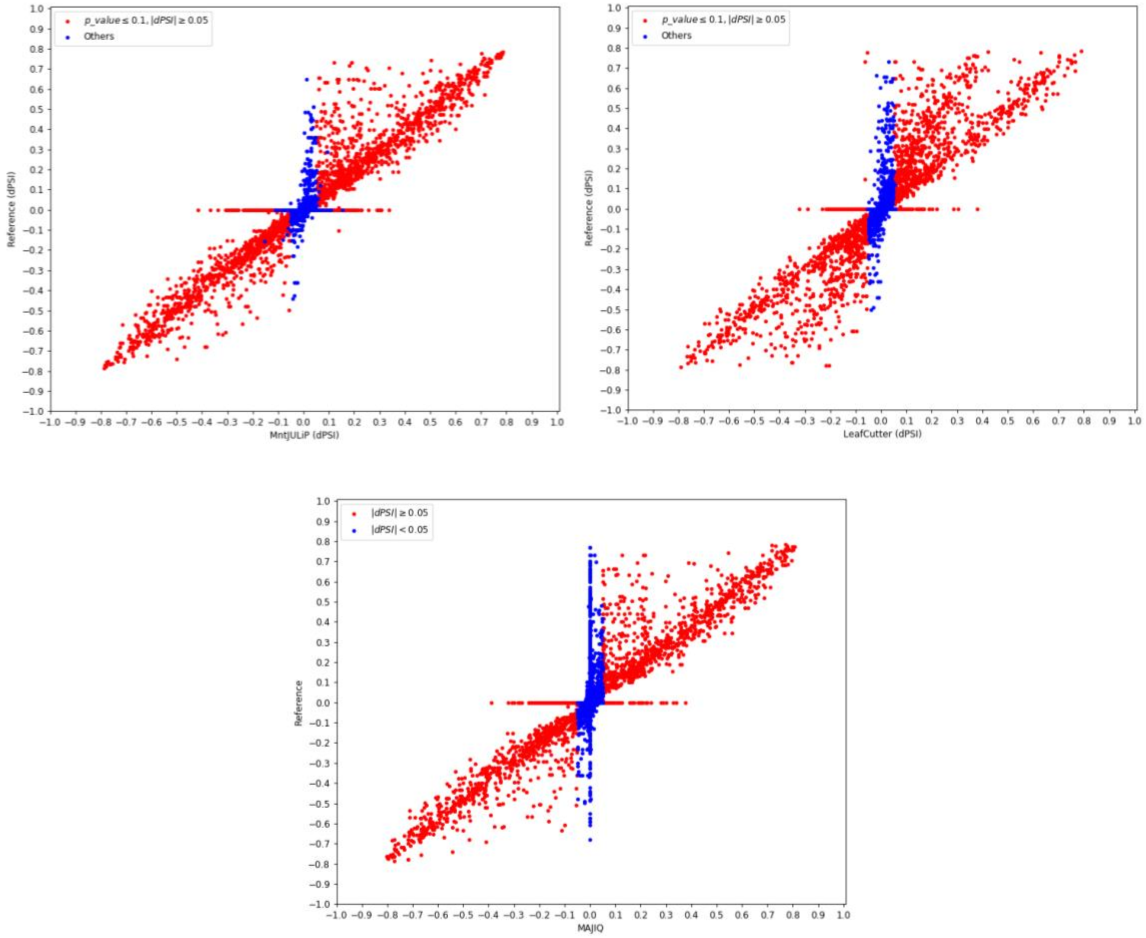

(B)

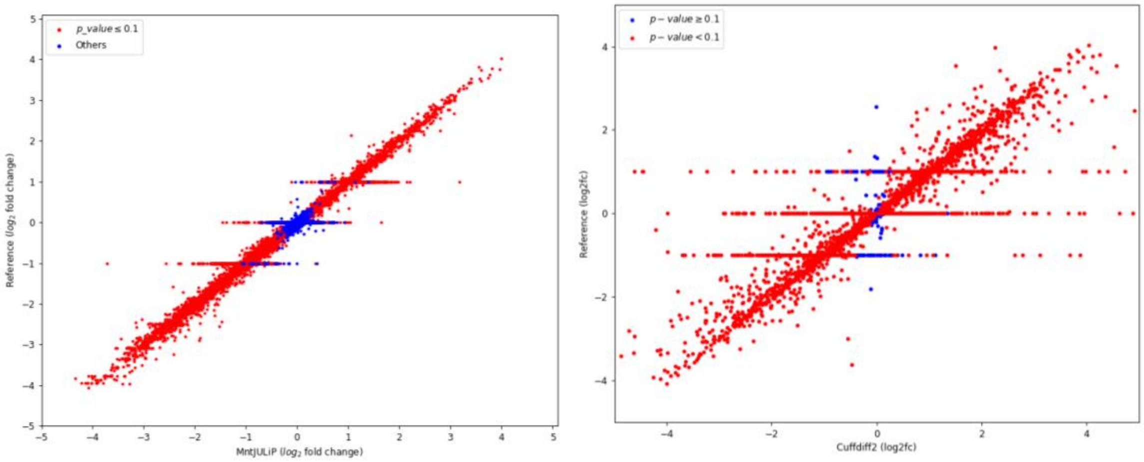


(C)

| CC | All | Diff. spliced |
|---|---|---|
| DSR | | |
| MntJULiP(DSR) | 0.935 | 0.952 |
| LeafCutter | 0.879 | 0.890 |
| MAJIQ | 0.847 | 0.860 |
| DSA | | |
| MntJULiP(DSA) | 0.991 | 0.995 |
| Cuffdiff2 | 0.848 | 0.884 |

**Figure S3.** Venn diagram of programs' predictions on the mouse hippocampus data set.

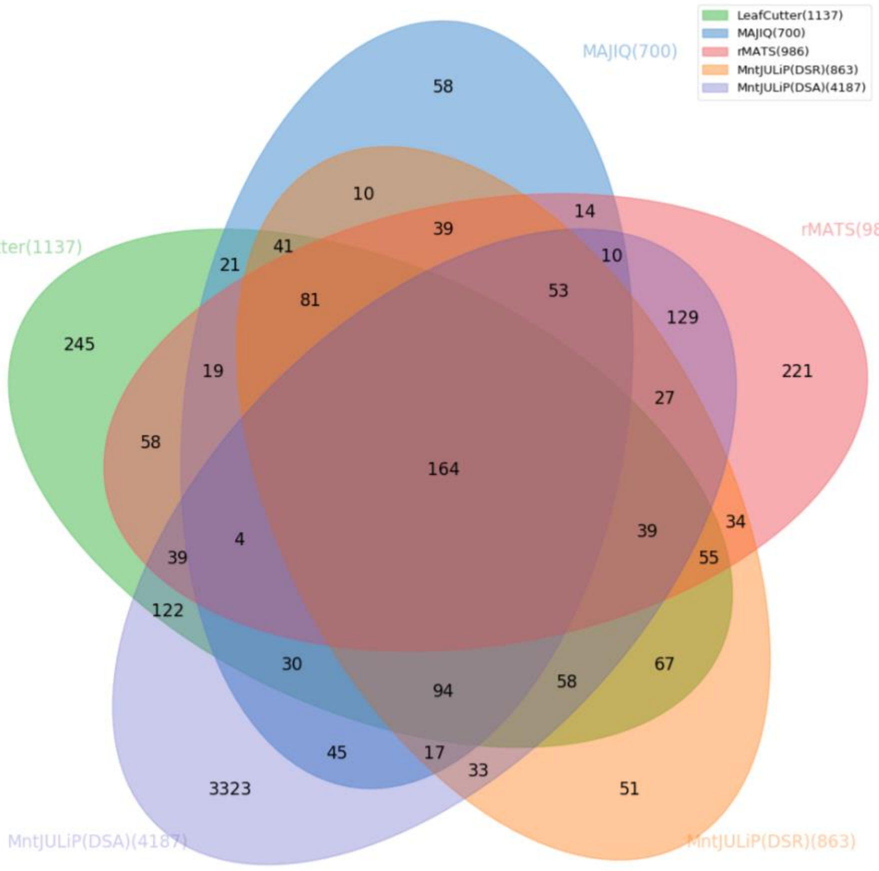

**Figure S4.** Examples of MntJULiP DSA predictions not identified by other tools (mouse hippocampus data set). (A) A mutually exclusive exon event at the *Pkm* gene locus is missed by DSR tools, but is identified by MntJULiP's DSA test via two of the flanking introns with significant changes in abundance. (B) The gene *Cwc22* harbors two overlapping and therefore mutually exclusively used introns, chr2:77881490-77903814 and chr2:77896578-77903796, with the former identified as significantly differentially abundant. (C) The gene *Cntf* shares its single intron (chr19:12,764,380-12,765,281) with the *Zfp91* gene. The intron is predicted by MntJulip DSA to undergo significant changes in abundance. The only *Zfp91* isoform containing this intron can be excluded based on the fact that one or more unique introns are not represented in the alignment data, therefore pointing to the gene *Cntf* being differentially expressed (*n.b.*, the differential expression of the gene *Cntf* was missed by DESeq2 due to its structure overlap with *Zfp91*). Note that the genes in (B) and (C) do not have *any* endpoint sharing introns, which is the required pattern for DSR-based methods, and therefore are *not* identifiable by any of the other tools.

(A)

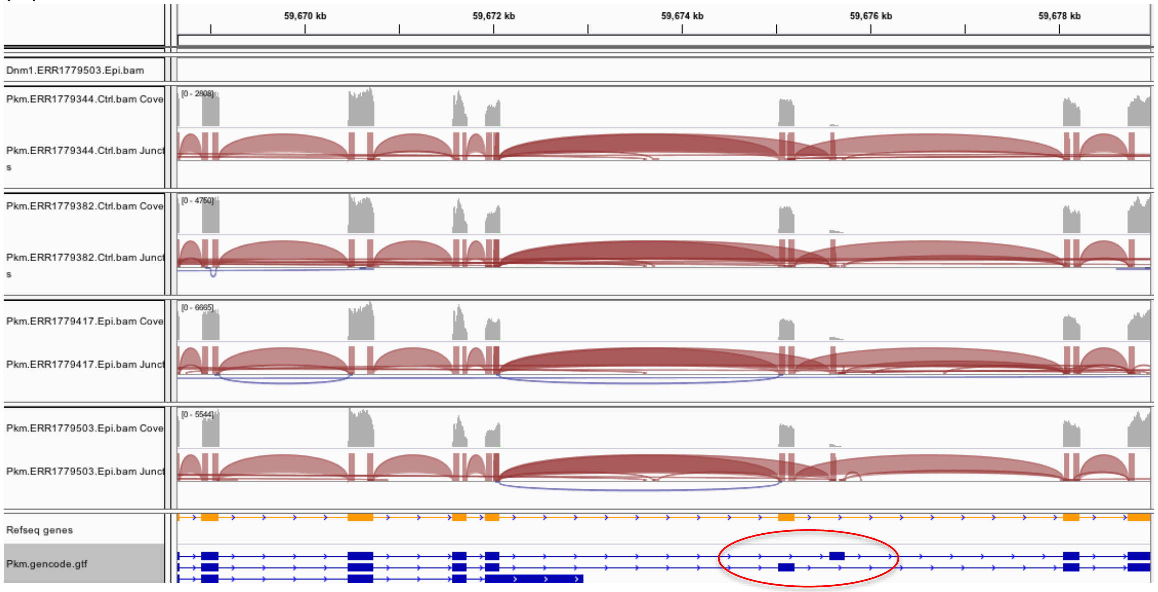

(B)

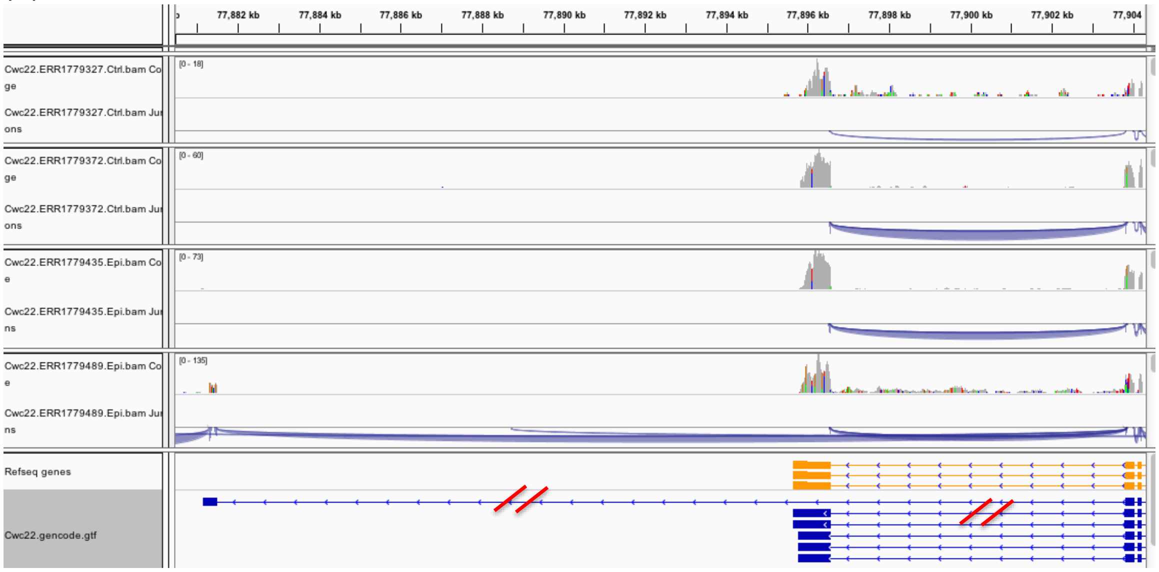
77,882 kb
77,884 kb
77,886 kb
77,888 kb
77,890 kb
77,892 kb
77,894 kb
77,896 kb
77,898 kb
77,900 kb
77,902 kb
77,904
Cwc22.ERR1779327.Ctrl.bam Coverage
[0 - 18]
Cwc22.ERR1779327.Ctrl.bam Junctions
Cwc22.ERR1779372.Ctrl.bam Coverage
[0 - 60]
Cwc22.ERR1779372.Ctrl.bam Junctions
Cwc22.ERR1779435.Epi.bam Coverage
[0 - 73]
Cwc22.ERR1779435.Epi.bam Junctions
Cwc22.ERR1779489.Epi.bam Coverage
[0 - 135]
Cwc22.ERR1779489.Epi.bam Junctions
Refseq genes
Cwc22.gencode.gtf

(C)

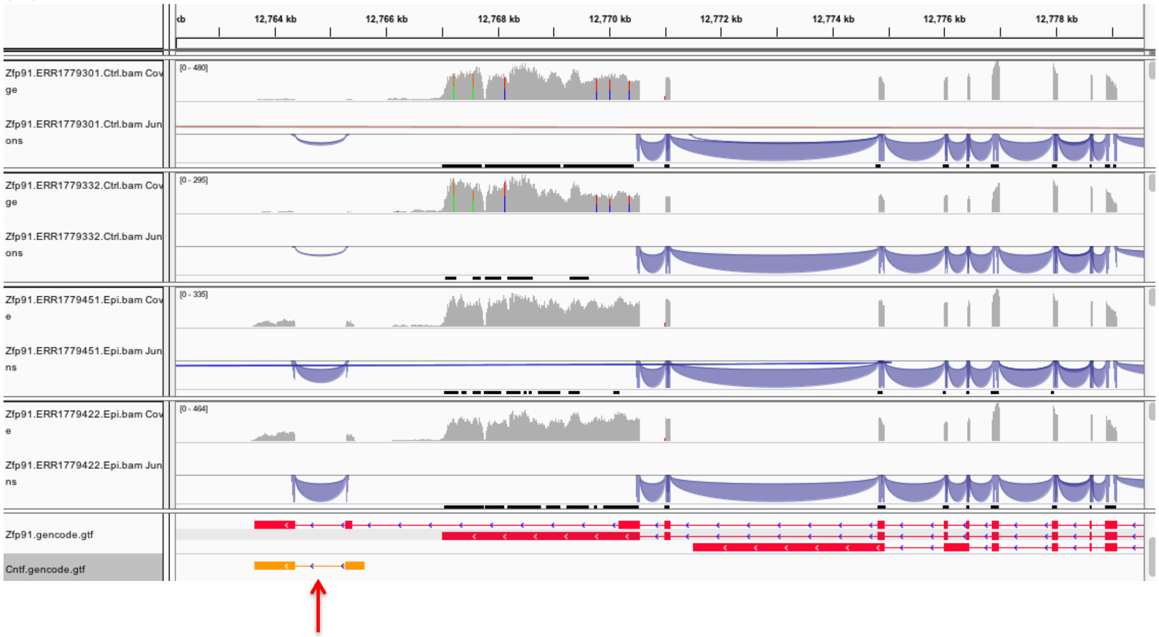
12,764 kb
12,766 kb
12,768 kb
12,770 kb
12,772 kb
12,774 kb
12,776 kb
12,778 kb
Zfp91.ERR1779301.Ctrl.bam Coverage
[0 - 480]
Zfp91.ERR1779301.Ctrl.bam Junctions
Zfp91.ERR1779332.Ctrl.bam Coverage
[0 - 295]
Zfp91.ERR1779332.Ctrl.bam Junctions
Zfp91.ERR1779451.Epi.bam Coverage
[0 - 335]
Zfp91.ERR1779451.Epi.bam Junctions
Zfp91.ERR1779422.Epi.bam Coverage
[0 - 464]
Zfp91.ERR1779422.Epi.bam Junctions
Zfp91.gencode.gtf
Cntf.gencode.gtf

**Figure S5.** Reproducibility plots for MntJULiP, LeafCutter and MAJIQ (DSR test) and MntJULiP (DSA test) on the mouse hippocampus data (A-D). Mouse hippocampus samples were divided randomly into two sets of 10x12 samples (healthy versus epileptic) each, and the per intron dPSI values (log2fc) predicted by each program are plotted between the two comparisons. Number of introns represented: 16,607 for MntJULiP (DSR), 30,738 for LeafCutter, 93,642 for MAJIQ, and 132,289 for MntJULiP (DSA). Correlation coefficients for the 4 comparisons are 0.579 for MntJULiP (DSR), 0.460 for LeafCutter, 0.577 for MAJIQ, and 0.665 for MntJULiP (DSA).

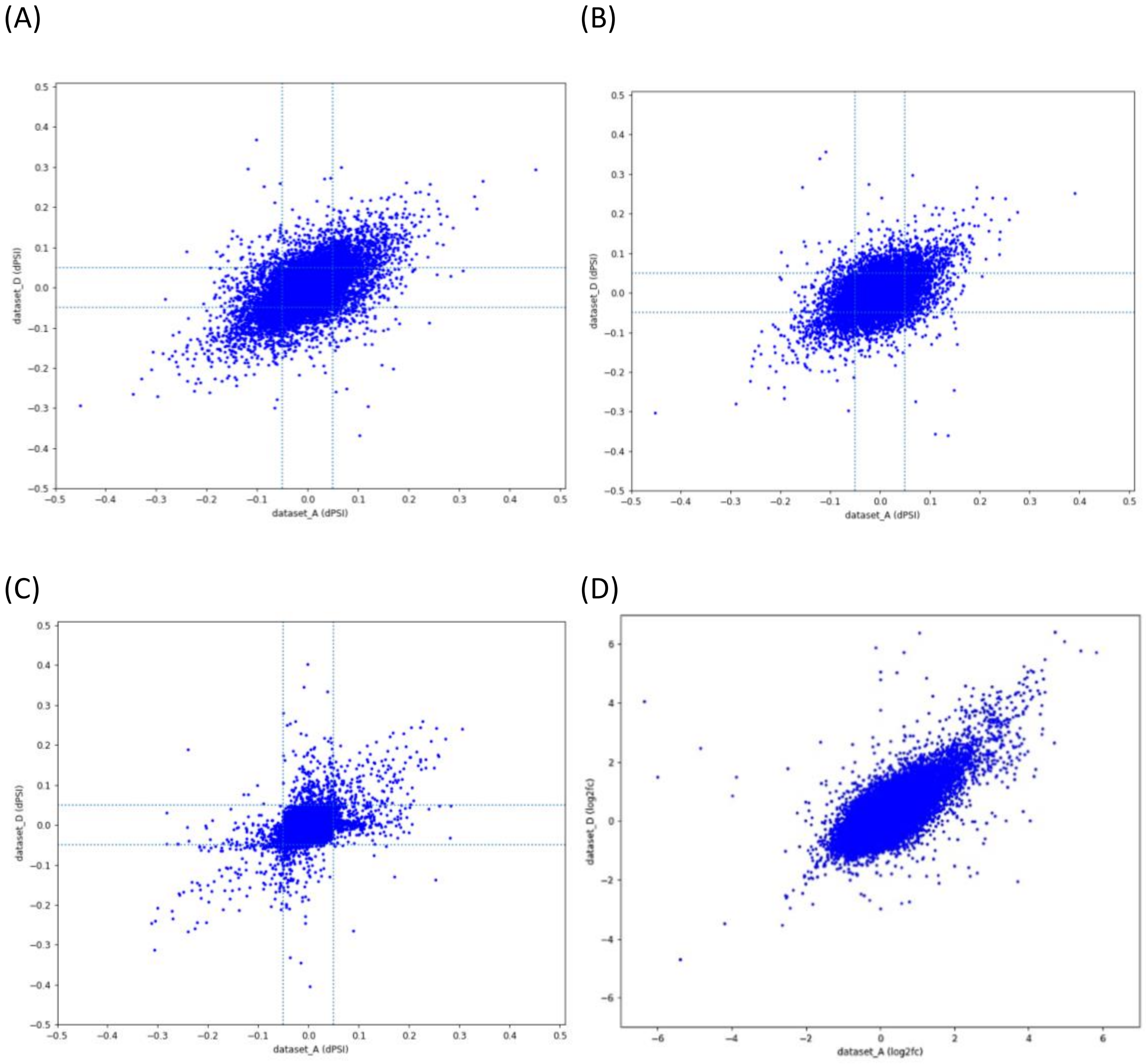

**Figure S6.** Multi-way versus all-against-all pairwise comparisons on GTEx tissue samples - gene sets. (A-B) Venn diagram of gene sets predicted by MntJULiP DSR (DSA) in comparisons of frontal cortex (105 samples), cortex (121 samples) and cerebellum (132 samples) RNA-seq collections. (C-D) Venn diagram of gene sets predicted by MntJULiP DSR (DSA) in comparisons of cortex (121 samples), cerebellum (132 samples) and lung (196 samples) RNA-seq collections. (A p-value cutoff of 0.05 was used for all comparisons.)

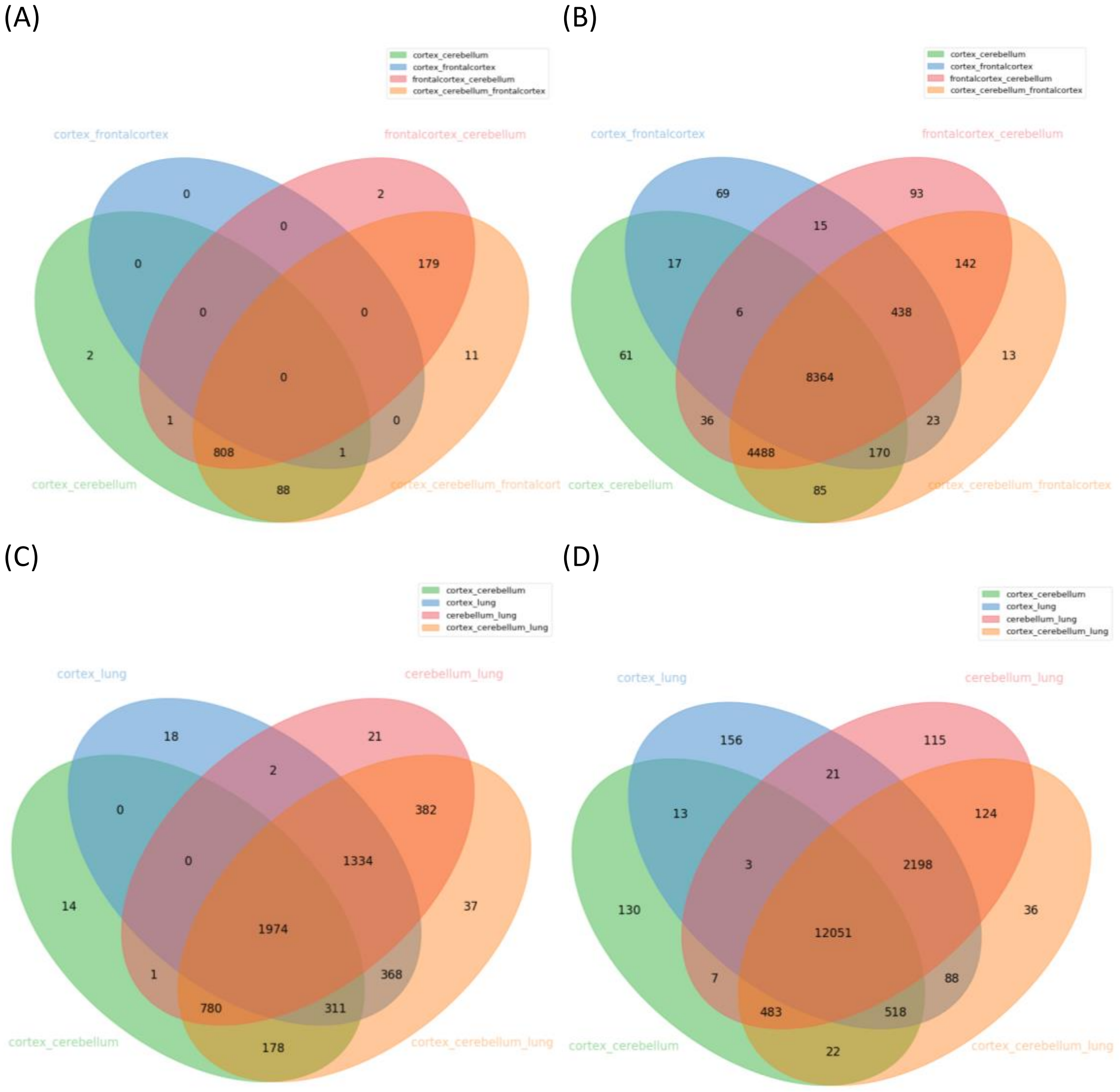

**Figure S7.** Multi-way versus all-against-all pairwise comparisons on GTEx tissue samples - heatmaps. (A) Heatmap of MntJULiP-identified DSR introns for the three-way and pairwise comparisons of frontal cortex, cortex and cerebellum collections. (B) Heatmap of MntJULiP-identified DSR introns in the three-way comparison of cortex, cerebellum and lung data collections. (C) Heatmap of MntJULiP-identified DSA introns for the three-way and pairwise comparisons of frontal cortex, cortex and cerebellum data collections. (D) Heatmap of MntJULiP-identified DSA introns in the three-way comparison of cortex, cerebellum and lung data collections. Note: Introns from genes with >30 reads across all samples, with dPSI>=0.2 (for DSR) and q-value<0.05, were plotted. For DSA, additionally, only the intron with the largest log 2 fold change was chosen to represent the gene.

(A)

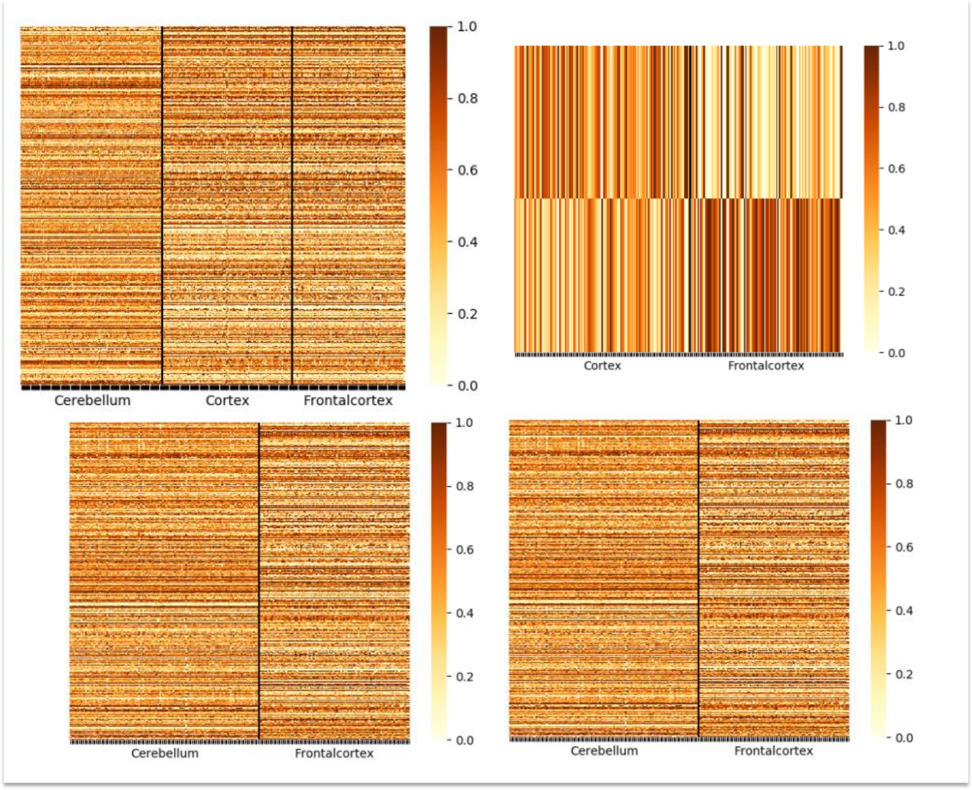


(B)

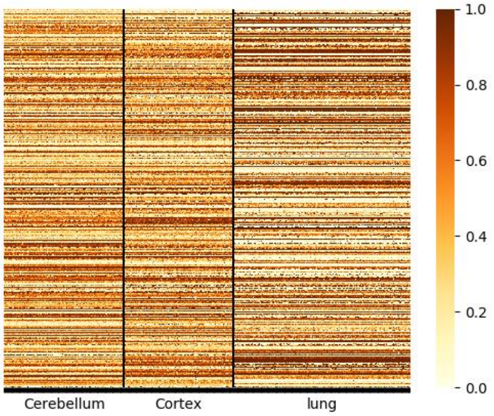

(C)

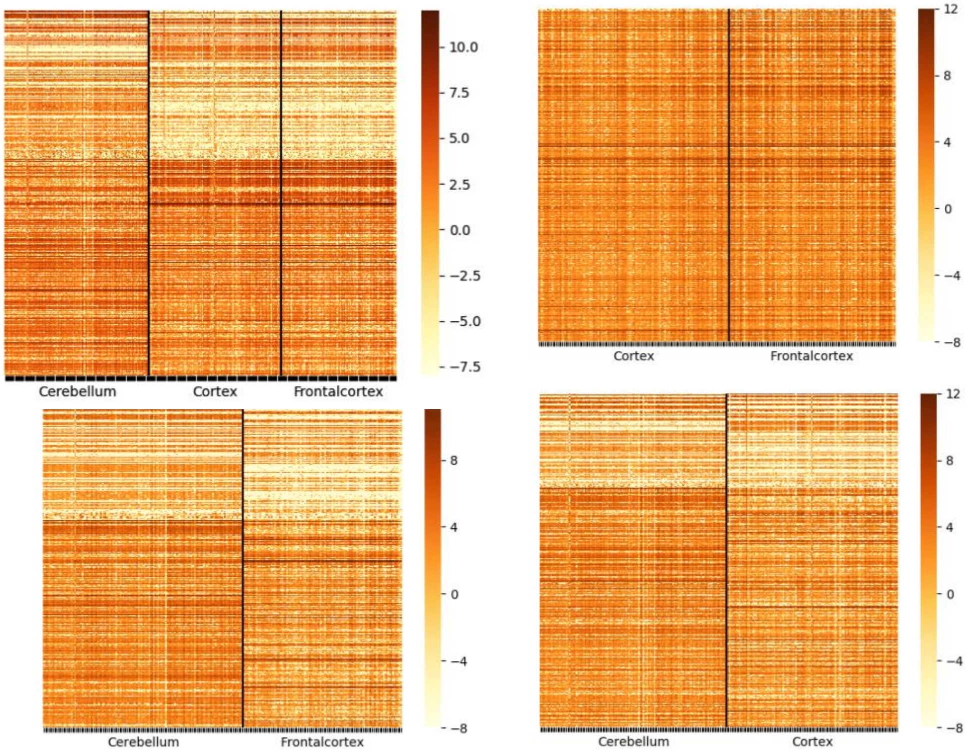
Cerebellum
Cortex
Frontalcortex
10.0
7.5
5.0
2.5
0.0
−2.5
−5.0
−7.5
Cortex
Frontalcortex
12
8
4
0
−4
−8
Cerebellum
Frontalcortex
8
4
0
−4
−8
Cerebellum
Cortex
12
8
4
0
−4
−8

(D)

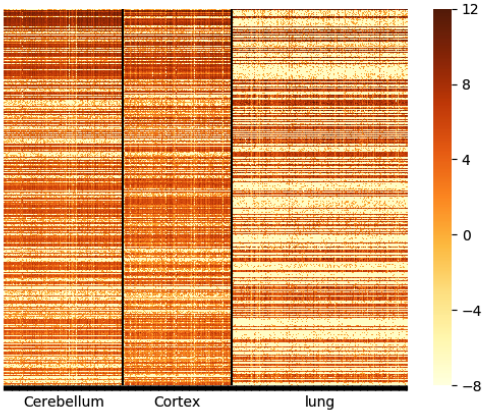
Cerebellum
Cortex
lung
12
8
4
0
−4
−8

**Figure S8.** Assessment of program predicted features for the time-series taste organoid data set, based on the assumption of continuity of feature space. The distribution of program-predicted features by number of comparisons is shown for three methods: i) union of MntJULiP predicted features from all (21 total) pairwise comparisons, ii) MntJULiP multi-way predicted features, and iii) union of LeafCutter predicted features, from all pairwise comparisons (21 total). For MntJULiP multi-way predictions, features were traced back to the pairwise comparisons in which they were reported (from i).

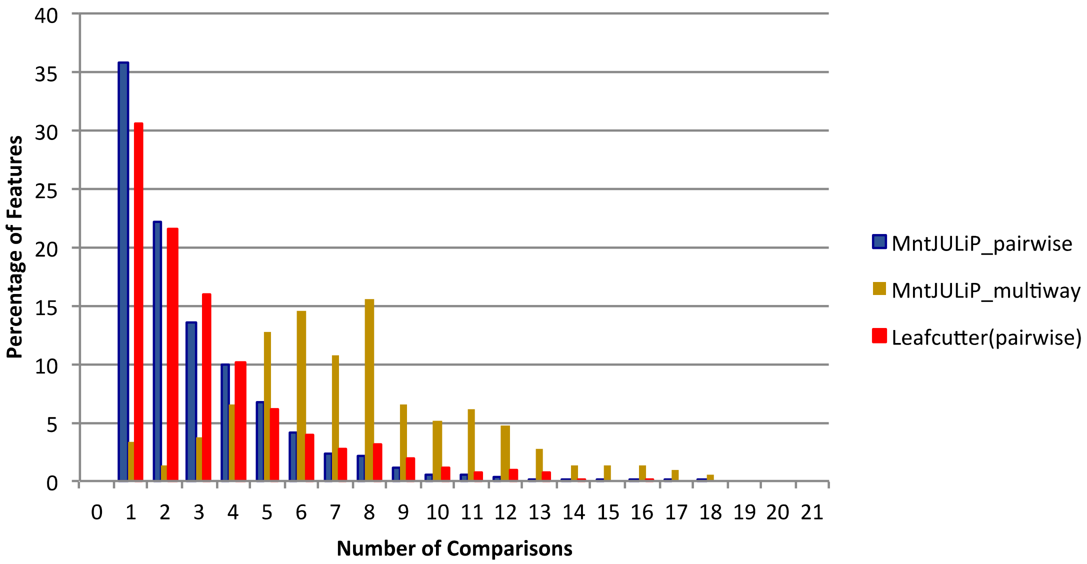

**Figure S9.** Heatmaps of differentially spliced features (introns) in the taste organoid data set. (A) MntJULiP DSR, features discovered via all pairwise versus multi-way comparison: (left) ‘all pairwise’ features, heatmap clustered by rows and columns; (center) ‘multi-way’ features, heatmap clustered by rows and columns; (right) ‘multi-way’ features, clustered by row (feature), samples ordered according to the time series. (B) Similarly, for MntJULiP DSA-predicted features. Features were filtered at p-value<0.05 and dPSI>=0.2 (for DSR). Grouping was performed using weighted hierarchical clustering with the Bray-Curtis metric.

(A)

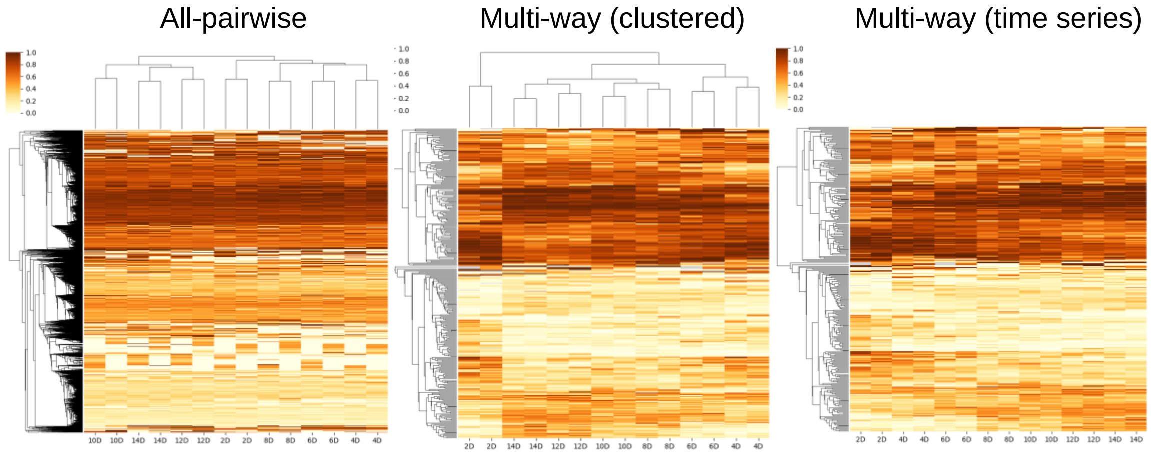


(B)

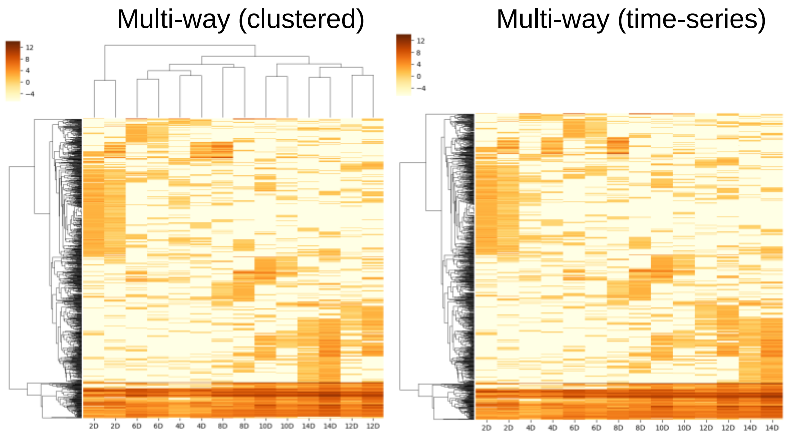
